# Bacterial Swarming-Guided Biomineralization Enables Pattern Formation in Engineered Living Materials

**DOI:** 10.64898/2026.05.05.722913

**Authors:** Koppisetty Viswa Chaithanya, Uttam Kumar, Karthik Pushpavanam

**Affiliations:** Chemical Engineering, Indian Institute of Technology, Gandhinagar, Gujarat, India, 382055; Institute of Physics, Academia Sinica, Taipei City, Taiwan, 115201

**Author notes:** To whom all correspondence must be addressed Karthik Pushpavanam, Ph.D., Department of Chemical Engineering, Indian Institute of Technology Gandhinagar, Gujarat, India, 382055, India.

**Keywords:** Swarming, Biomineralization, Engineered Living Materials, Patterning, Calcium Hydroxyapatite

## Abstract

Engineered living materials (ELMs) harness the adaptive and self-replicating capabilities of biological systems to create functional materials for sensing, catalysis, and biomineralization. While most ELM strategies rely on static microbial assemblies, the role of bacterial motility in structuring living materials remains unexplored. Here, for the first time, we demonstrate how swarming motility in *Escherichia coli* MG1655 can be induced to guide spatio-temporally organized calcium phosphate mineralization. The mineralized calcium phosphate is characterized by scanning electron microscopy and elemental analysis. By systematically varying phosphate sources and their concentrations in calcium-rich media, we observe the emergence of regularly spaced concentric mineralized patterns. The previously undocumented observation of the concentric patterns was rationalized through a continuum model that captures the spatiotemporal coupling between swarm expansion and mineral deposition. The model shows that this coupling can generate recurrent front arrest and restart, leading to concentric ring formation. Finally, we show that altering the phosphate species results in distinct mineral morphologies. Together, this work establishes a novel framework for integrating bacterial swarming with biomineralization, enabling dynamic and programmable pattern formation in ELMs.

## INTRODUCTION

The synthesis of functional inorganic materials conventionally relies on top-down fabrication or bottom-up chemical strategies, including sol-gel processing, high-temperature sintering, and chemical precipitation, providing precise control over morphology and crystallinity^1–5^. However, these approaches are resource-intensive and operate under harsh conditions^6,7^. Biomineralization mediated by a diverse palette of biological macromolecules- most notably proteins, peptides, and polysaccharides present a compelling alternative to fabricating inorganic materials under ambient conditions^8–10^. These biomolecules orchestrate nucleation, growth, and assembly of inorganic phases with exquisite spatiotemporal control^11^. Despite these advances, these approaches are inherently limited in their scalability and regenerative capacity. Isolated biomolecules cannot self-replenish or adapt dynamically to changing conditions. This constraint has motivated a paradigm shift toward engineered living materials (ELMs), platforms in which biological systems serve as active, self-sustaining factories for inorganic material synthesis^7,12,13^.

A particularly promising avenue within the ELM field is the use of microbial biomineralization to generate living composites^14–16^. ELMs offer a powerful platform for biomineralization by leveraging the intrinsic biological capabilities, including assembly, environmental responsiveness, and self-repair, as features inaccessible to conventional synthetic approaches^17,18^. By embedding living microbes as active components, ELMs facilitate continuous coupling between cellular metabolism and material synthesis, allowing dynamic control over mineralization processes. Microorganisms regulate their local physicochemical environment through ion binding, metabolic pH modulation, and enzymatic activity, thereby promoting localized supersaturation and acting as nucleation centers for mineral deposition^13,19^. Direct experimental observations further confirm that bacterial cell surfaces bind Ca^2+^ and serve as nucleation sites for crystal growth during biomineralization processes^20^.

Microbially induced calcium biomineralization has been widely demonstrated through processes such as microbially induced calcium carbonate precipitation (MICP) and microbially induced phosphate precipitation (MIPP), where cellular metabolism drives local supersaturation and mineral formation^14,26–28^. While recent studies employing engineered strains have demonstrated the ability to modulate mineral formation pathways, these approaches remain largely constrained to static or biofilm-based systems and lack dynamic spatiotemporal control over mineral architecture^12,14,21^. These ELM systems largely depend on preformed scaffolds or immobilized cells, further restricting their diversity in applicability ^28,40^.

Swarming motility, in contrast, provides a dynamic mode of collective organization, although it is often considered to exhibit strain-dependent patterning behavior^29–31^. Challenging this paradigm, we demonstrate that the native swarming patterns of *Escherichia coli* can be reprogrammed through environmental modulation to mimic concentric ring-like expansion patterns typically associated with *Proteus mirabilis*, without genetic alteration. Furthermore, by controlling phosphate availability under these swarming conditions, we show that mesoscale mineral deposition can be directed, resulting in distinct spatial organization of calcium phosphate (hydroxyapatite-like) structures. These findings establish a direct link between environmentally tunable collective bacterial behavior and mineral morphology, providing a route to dynamically controlled biomineralization that extends beyond the limitations of existing static ELM and biofilm-based systems.

## MATERIALS AND METHODS

### Materials

We sourced the materials from the following suppliers.

### SRL Pvt. Ltd

Calcium Chloride (cat# 70650), Sodium Phosphate Dibasic (cat# 66273), Sodium Phosphate Monobasic (cat# 59443), Potassium Phosphate Dibasic (cat# 27387), Potassium Dihydrogen Orthophosphate (cat# 54358). **HiMedia:** Nutrient Broth No. 3 (cat# M1093), Agar Agar, Type I (cat# GRM666). Tokyo Chemical Industry (TCI) Co. Ltd: D-(+)-Glucose (cat# G0048). **Tarsons Products Ltd:** 6-well cell culture plates (cat# 980010). **Sigma-Aldrich:** β-Glycerophosphate (cat# 35675).

### Bacterial growth conditions

The *Escherichia coli* MG1655 strain (MTCC No. 1586) was used in this study. All bacterial cultures were propagated on Nutrient Broth 03. For swarming experiments, nutrient agar plates containing 0.5% agar were used. The specific concentrations of phosphate and calcium chloride were achieved by adding the respective 1 M stock solutions to molten nutrient agar at 45 °C. Glucose-containing media were prepared by adding the required amount of glucose and the components before sterilization. A single colony was inoculated into nutrient broth 03 and incubated at 37 °C overnight; the resulting saturated culture was used for the swarming experiments.

### Phase-contrast Microscopy

The Nikon Eclipse Ts2R inverted microscope was used for all of our experiments. NIS-Elements BR version 6.20.01 64-bit software accompanying the machine was used to observe and acquire the image of the swarming bacteria in 6-well plates, directly mounted onto the stage in the phase contrast (Ph1) setting. The measurements performed on the images were aided by the same software.

### Characterization through scanning electron microscopy (SEM)

The swarm area, along with nutrient agar, was dried onto aluminum foil, and an appropriate portion was analyzed by SEM (JSM-7600F, JEOL). Each surface was made conductive by applying a platinum coating for 1 min, and observations were performed in Low Energy Imaging (LEI) mode at 5 kV. Additionally, Energy-Dispersive Spectroscopy (EDS) was performed to analyze the elemental composition of selected areas of the sample.

## RESULTS AND DISCUSSION

We first examined whether the phosphate source could influence the swarming behavior of *E. Coli* MG1655 compared to its growth on Nutrient Broth 03 (control) (**Figure 1A**), as changes in motility could modulate mineral formation. To test this, MG1655 cells were spotted on swarming agar supplemented with different phosphate sources. These included β-glycerophosphate, sodium and potassium monobasic phosphates (NaH_2_PO_4_, and KH_2_PO_4_), and dibasic phosphates (Na_2_HPO_4_, and K_2_HPO_4_), each at 10 mM. The swarming patterns observed across all phosphate-supplemented conditions were comparable to those on the control plate (Nutrient Broth 03) (**Figure 1B-F**). This indicates that the tested phosphate sources do not significantly affect MG1655 swarming under the conditions tested. In contrast, the concentration of CaCl_2_ as low as 5 mM severely affected the swarming pattern and restricted bacterial motility, which was further hindered at 10 and 20 mM (**Figure 1G-I**). We posit that Ca^2+^ ions could directly interact with and perturb the cell envelope and ionic balance, whereas phosphate is a regulated nutrient that is assimilated and buffered by the cell, resulting in minimal toxicity under comparable conditions^44^.

**Figure 1.**
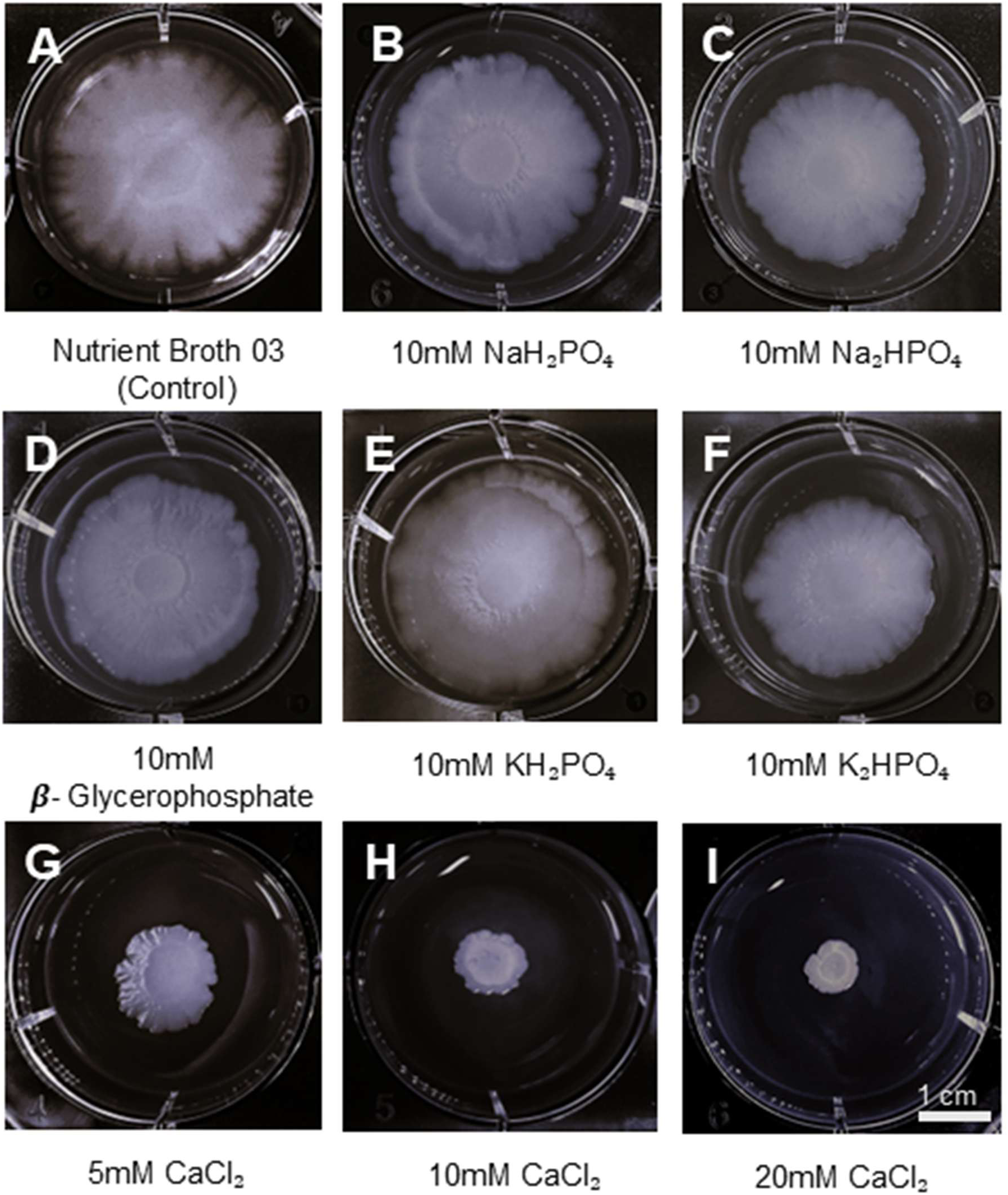
Effect of phosphate and calcium on bacterial swarming. Swarming pattern of bacteria observed after incubation for 4 days at 37℃ grown in **(A)** Nutrient broth 03 (Control), **(B)** 10mM NaH_2_PO_4_, **(C)** 10mM Na_2_HPO_4_, **(D)** 10mM ***β***- Glycerophosphate, **(E)** 10mM KH₂PO₄, **(F)** 10mM K_2_HPO_4_, **(G)** 5mM CaCl_2_, **(H)** 10mM CaCl_2_ and, **(I)** 20mM CaCl_2_.

Having established the compatibility of phosphate with swarming, we next sought to identify a calcium-phosphate combination capable of supporting bacterially mediated mineralization without abiotic precipitation confounding the results. When dibasic phosphate salts (Na_2_HPO_4_ or K_2_HPO_4_) were combined with CaCl_2_ in the medium, cloudy precipitates formed spontaneously, whereas no such precipitates were observed with monobasic salts (**Figure S1**). This abiotic precipitation is consistent with the inherent basicity of dibasic salts, which elevates local pH and drives spontaneous calcium phosphate precipitation^45^. To isolate bacterially mediated mineralization from this abiotic background, we selected 10 mM Na_2_HPO_4_ and 10 mM CaCl_2_ as the baseline mineralizing medium for subsequent experiments. When *E. Coli* MG1655 was spotted onto this mineralizing medium, swarming produced a white thin film across the colony’s surface (**Figure 2A**). This morphology was visibly distinct from the featureless control swarm (**Figure 1A**). Strikingly, well-defined concentric disc-like structures developed radially in concert with the expanding swarm front (**Figure 2B**).

**Figure 2.**
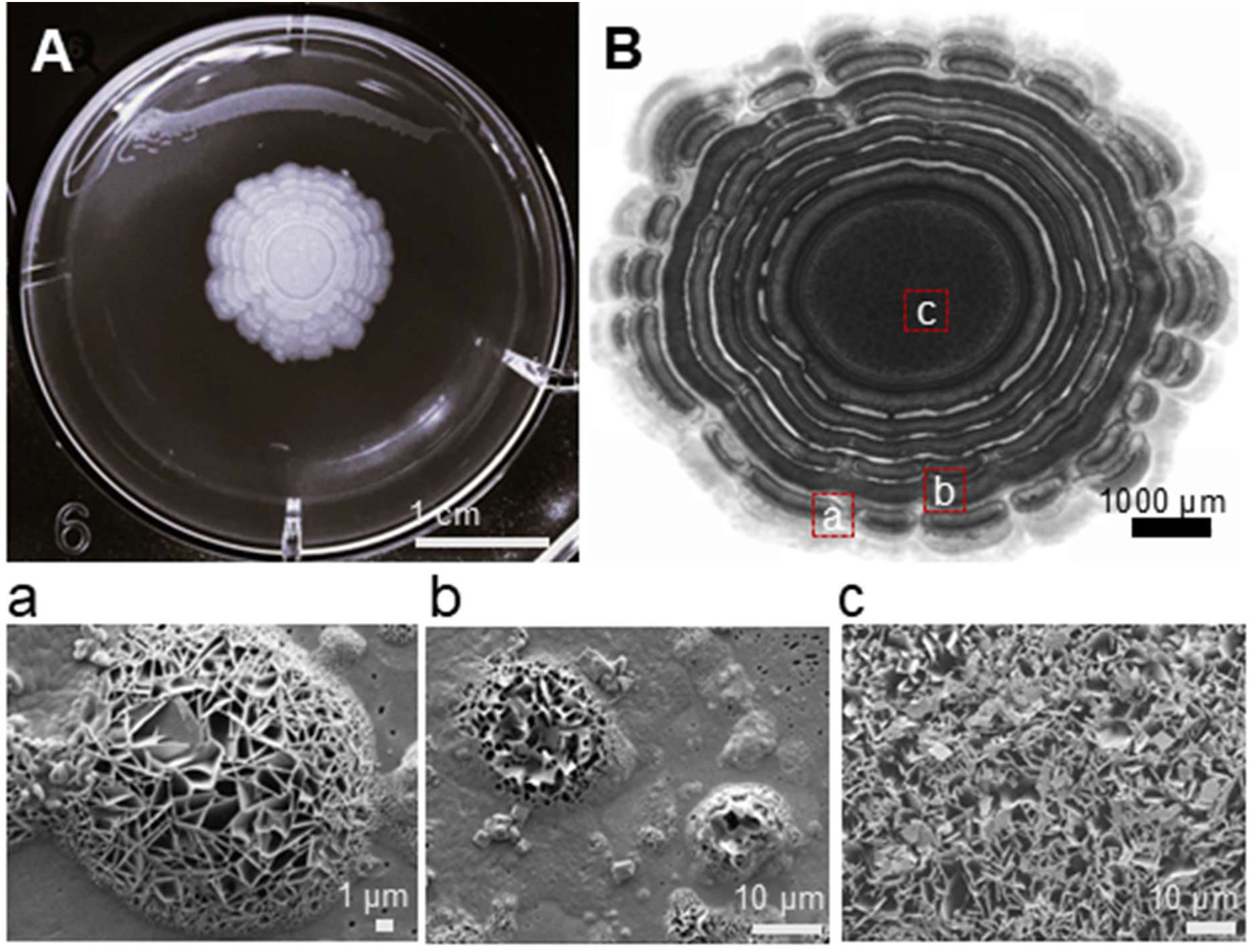
Concentric mineralization of calcium chloride during bacterial swarming. **(A)** The swarming pattern of bacteria observed after incubation for 4 days at 37℃, grown in Nutrient broth supplemented with 10 mM CaCl_2_ and 10 mM NaH_2_PO_4_, and **(B)** Phase-contrast micrograph of the swarm, with marked alphabets (a-c) indicating approximate zones corresponding to mineralized calcium phosphate under a scanning electron microscope.

To characterize the spatiotemporal dynamics of this phenomenon, we collected optical micrographs every 24 hours. New disc-like structures consistently appeared near the swarm front, while older rings remained immobile (**Figure S2A–C**), indicating that disc formation is a continuous process coupled to the outward swarm progression. Newly formed bands appeared optically light initially, becoming progressively thicker and more opaque over time. In the course of expansion, mineralizing swarm fronts showcase particulate mineral inclusions in the liquid film (**Supplementary** V**ideo S1**). We hypothesize that this likely due to the presence of mineralized calcium interspersed between the motile bacteria. The emergence of these previously unreported concentric patterns in the *E.coli* prompted us to determine whether they represented organized mineral deposition.

To determine the chemical identity and surface morphology of the thin film, we employed scanning electron microscopy (SEM) coupled with energy-dispersive X-ray spectroscopy (EDS). SEM imaging revealed that distinct zones across the swarm exhibited reproducible mineralization patterns (**Figure 2C**). EDS analysis of swarms grown under equimolar CaCl_2_ (10 mM) and NaH_2_PO_4_ (10 mM) yielded a mean Ca:P molar ratio of 1.679 ± 0.04 (**Figure S3**). This value closely approximates the stoichiometric Ca:P ratio of 1.6 characteristic of calcium hydroxyapatite (Ca_10_(PO_4_)_6_(OH)_2_), identifying HAp as the primary mineral phase deposited during swarming.

To validate that mineral deposition requires the combined presence of bacteria, calcium, and phosphate, we examined control swarms grown in NB03 alone, NaH_2_PO_4_ alone, and CaCl_2_ alone by SEM. In all three cases, swarms appeared as flat, planar cell layers with no mineralization (**Figure S4A–C**). EDS of the NB03 control detected only trace calcium attributable to media components (**Figure S5**). Notably, EDS of the calcium-supplemented swarm grown without phosphate also yielded lower calcium signal than mineralized swarms (2.5 % vs 31.1 %) (**Figure S6**), consistent with reversible, non-precipitative calcium binding by bacterial cell surfaces^27^. Collectively, these confirm that bacterially driven HAp deposition is contingent on the simultaneous availability of both calcium and the inorganic phosphate.

With the chemical identity of the mineral deposits established, we next investigated how key experimental variables like initial cell density and calcium concentration influence the spatial organization of biomineralization. To examine the effect of cell density, we spotted inoculum volumes ranging from 1.25 to 10 µl onto mineralizing media. As anticipated increasing inoculum volume expanded the total circumference of the mineralized swarm (**Figure S7**). This observation is consistent with accelerated mineral deposition at higher cell densities, where greater metabolic activity and phosphate processing capacity are likely to drive faster HAp formation. We then examined the effect of calcium concentration on mineralization patterning by varying CaCl_2_ from 5 to 20 mM in the presence of 10 mM NaH_2_PO_4_. Increasing calcium concentration progressively reduced the spacing between consecutive concentric bands, yielding more densely packed ring structures at higher calcium levels (**Figure S8**). Importantly, HAp morphology was conserved across all tested calcium concentrations, with more compact HAp formations at higher calcium concentrations (**Figure S8**), demonstrating the robustness of the bacterially mediated mineralization process. Taken together, these results show that both cell density and calcium availability serve as tunable parameters for modulating the spatial frequency and extent of HAp deposition.

The spontaneous precipitation observed with dibasic phosphate salts implicated pH as a potentially important regulatory variable in this system. To further probe pH as a parameter influencing mineralization, we supplemented the media with glucose. Glucose is a fermentable carbon source that is pH-neutral in isolation, but whose bacterial catabolism generates acidic byproducts, progressively lowering environmental pH during growth ^46^. Under standard conditions, bacterial growth on NB03 and mineralizing conditions increased media pH (**Figure S9**). As expected, the glucose supplementation caused a measurable decrease in pH (**Figure S9**). This acidic microenvironment generated by glucose metabolism substantially impaired swarming motility in glucose added NB03, 10mM CaCl_2_ and 10mM NH_2_PO_4_ conditions (**Figure S10A-C**). Correspondingly, in the mineralizing media, glucose generated an acidic environment that simultaneously disrupted swarming and abolished concentric mineralization (**Figure S10D-F**). SEM imaging further revealed elongated cellular morphology under glucose-rich conditions relative to standard growth on NB03 (**Figure S11**). This observation aligns with previous studies documenting that bacterial cell size is strongly influenced by nutrient availability, with *E.coli* exhibiting dynamic cell length relative to nutrient abundance^47^. These findings collectively demonstrate that acidic pH functions as a negative regulator of both swarming motility and calcium phosphate biomineralization under the conditions tested and suggest that the alkalinization accompanying normal swarming is conducive for HAp mineralization.

Having established the conditions under which bacteria reliably produce HAp, we asked whether deliberate selection of the phosphate source could be used to tune the microstructure of the resulting mineral, an important consideration for potential applications in materials design. We compared KH_2_PO_4_ (inorganic monobasic phosphate) and β-glycerophosphate (organic phosphate ester) as phosphate sources in the presence of 10 mM CaCl_2_ and characterized the resulting mineralization by SEM (**Figure 3**). Both conditions supported thin-film mineral deposition within the swarm. Scanning electron micrographs revealed markedly distinct microarchitectures: KH_2_PO_4_ produced ridge-like surface patterns (**Figure 3B**) comparable to those observed with NaH_2_PO_4_, while β-glycerophosphate gave rise to dense, labyrinthine structures with a conspicuously more compact morphology (**Figure 3D**). Concentric banding was more prominent under KH_2_PO_4_ than under β-glycerophosphate conditions, suggesting that the rate and mode of phosphate delivery also influence mesoscale patterning. EDS analysis yielded Ca:P ratios of 1.26 and 1.71 for KH_2_PO_4_ and β-glycerophosphate-induced mineralization, respectively (**Figure S12 and S13**).

**Figure 3.**
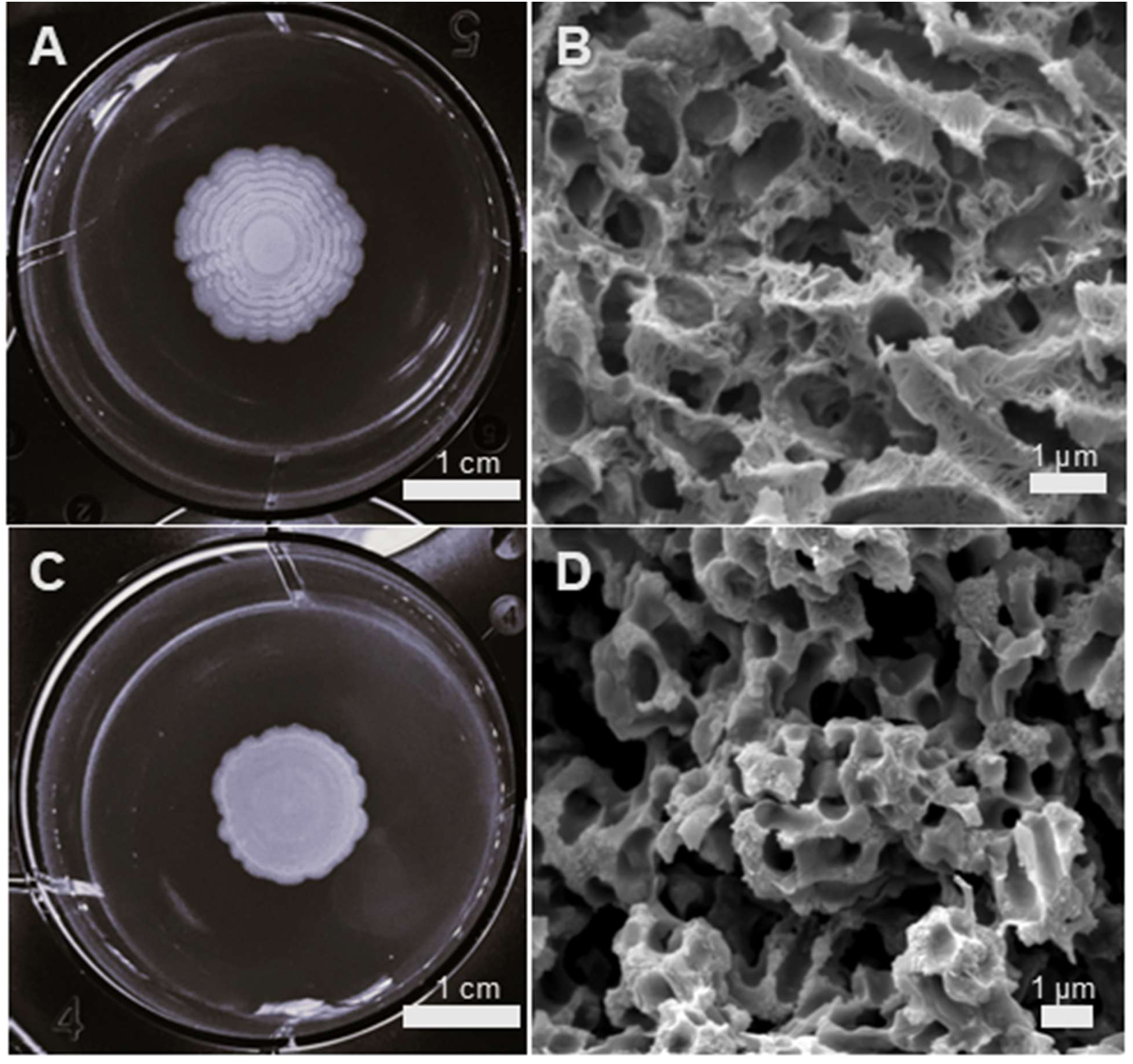
Bio-mineralized calcium hydroxyapatite by modulating phosphate source. SEM images showing precipitate characteristics and geometry can be controlled by modifying the phosphate source to **(A, B)** 10 mM KH_2_PO_4_ and **(C, D)** 10 mM β-glycerophosphate in a calcium-rich (10 mM CaCl_2_) environment.

We propose that these structural and compositional differences reflect the distinct phosphate release kinetics of the two sources. As an inorganic salt, NaH_2_PO_4_ and KH_2_PO_4_ dissociates freely in solution, providing immediately bioavailable phosphate that diffuses readily throughout the swarm, enabling rapid and spatially uniform mineralization. β-glycerophosphate, by contrast, is an organic phosphate ester that must first undergo enzymatic hydrolysis by bacterial phosphatases before inorganic phosphate is released. This gradual, enzymatically gated phosphate liberation likely produces localized phosphate microenvironments that favor the formation of denser, more tortuous mineral architectures. Collectively, these results demonstrate that the bacterially deposited HAp can be rationally controlled by selecting the appropriate phosphate precursor, offering a potential strategy for tailoring mineral properties via active matter driven synthesis.

The experimentally observed concentric banding raises a fundamental mechanistic question: what drives the repeated arrest and restart of the swarming front that gives rise to discrete mineral rings? To address this, we developed a continuum model of coupled bacterial swarming and calcium phosphate mineralization. Our framework is inspired by the terrace-forming swarming model of Ayati^42^ which proposes that coordinated swarming initiates only once local bacterial density exceeds a critical threshold, and that the front advances through successive arrest-and-restart cycles. We extend this framework to incorporate local calcium-phosphate supersaturation and the mechanical feedback of deposited mineral on bacterial motility.

The model is formulated in radial coordinates, where *r* denotes the distance from the inoculation point and t denotes time. Three coupled fields are tracked: *P*(*r*, *t*), the effective bacterial swarming activity at the colony front; *A*(*r*, *t*), the local activation state representing calcium-phosphate supersaturation; and *I*(*r*, *t*), the mineralization-related inhibition that suppresses motility following mineral deposition. The governing equations of the reduced model are

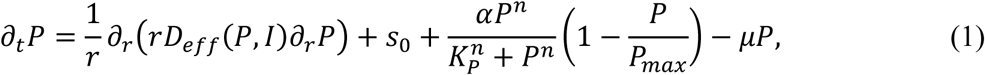

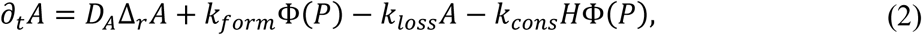

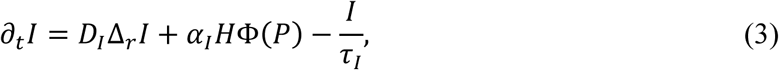

with,

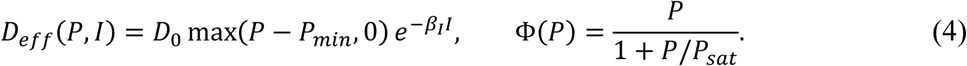

The equation for *P*(*r*, *t*) captures two defining features of ring-forming swarming. First, coordinated motion is only possible above a density threshold and is suppressed by local mineral deposition, as encoded in the effective motility *D_eff_*(*P*, *I*). Second, once swarming initiates, it is self-reinforcing via cooperative density-dependent activation, represented by the Hill-type term.

The equations for *A*(*r*, *t*) and *I*(*r*, *t*) are introduced phenomenologically to model the activation-inhibition cycle underlying repeated ring deposition. The activation state *A* builds up during swarming activity and is depleted as the front passes through a site and during a deposition event.

The local activation state *A* of a site builds up through *k_form_Φ*(*P*) and decays passively toward zero at rate *k_loss_* in the absence of firing. During an ACTIVE event (*H* = 1), it is depleted by *k_cons_HΦ*(*P*), causing *A* to fall from *A_on_* to *A_off_* and ends the firing event. Biologically, it represents local calcium-phosphate supersaturation that primes a site for mineralization. The inhibition *I* is produced during ACTIVE phase at rate *α_I_HΦ*(*P*) and relaxes afterward through −*I*/*τ_I_*. It is therefore generated only when *H* = 1, linking inhibitor production to the mineralization event rather than to constitutive bacterial activity. It spreads locally by diffusion at rate *D_I_* and decays at rate *I*/*τ_I_* once the deposition event ends. Biologically, it represents the mechanical obstruction imposed by deposited mineral on local motility.

To capture the irreversible nature of mineral deposition, each spatial site is assigned one of three discrete states: READY, ACTIVE, or REFRACTORY. A site transitions from READY to ACTIVE only when *A*(*r*, *t*) exceeds the threshold *A_on_*, local swarming activity *P*(*r*, *t*) exceeds *P_gate_*, and inhibition *I*(*r*, *t*) remains below *I_gate_*. The binary gate *H*(*r*, *t*) ∈ {0,1} equals 1 only at ACTIVE sites. Once a site transitions to REFRACTORY, it cannot be reactivated, since the deposited mineral constitutes a permanent local obstruction. This irreversible refractory memory is a key feature of the model, as it prevents re-firing at previously mineralized locations. After each deposition event, the inhibitor relaxes during the refractory phase, and the associated recovery time *T_off_* determines the interval before the front can resume motion and trigger the next ring-forming event. The threshold parameters *A_on_*, *A_off_*, *P_gate_*, and *I_gate_* are constrained by the steady states. Their derivation and consistency conditions are given in the Threshold parameter calculation in the Supplementary Section. All parameter values are listed in **Table S1**.

Numerical simulation of Eqs. (1)–(4) across biologically reasonable parameter values confirms that the proposed mechanism is sufficient to generate concentric ring patterns. For representative parameters *D_o_* = 1.0, *A_on_* = 0.70, *D_I_* = 0.010, and *D_A_* = 0, the model produces eight concentric rings with a mean spacing of *λ̄* = 4.71 model units (**Figure 4A**). To assess the robustness of ring formation, we performed a parameter space survey over a 3×3×3 grid spanning *D_o_* ∈ {0.70, 1.00, 1.30}, *A_on_* ∈ {0.55, 0.70, 0.85}, and *D_I_* ∈ {0.005, 0.010, 0.020} (**Figure 4B**). Ring formation was confined to a coherent region of parameter space centered on *A_on_* = 0.70 and *D_I_* ≤ 0.010. Outside this regime, the colony either advanced continuously without arrest or stalled permanently at the boundary. Notably, varying D₀ altered ring spacing without determining whether rings formed at all, indicating that the ring-forming instability is governed by chemical timescales, specifically the activation-inhibition cycle, rather than by swarming speed.

**Figure 4.**
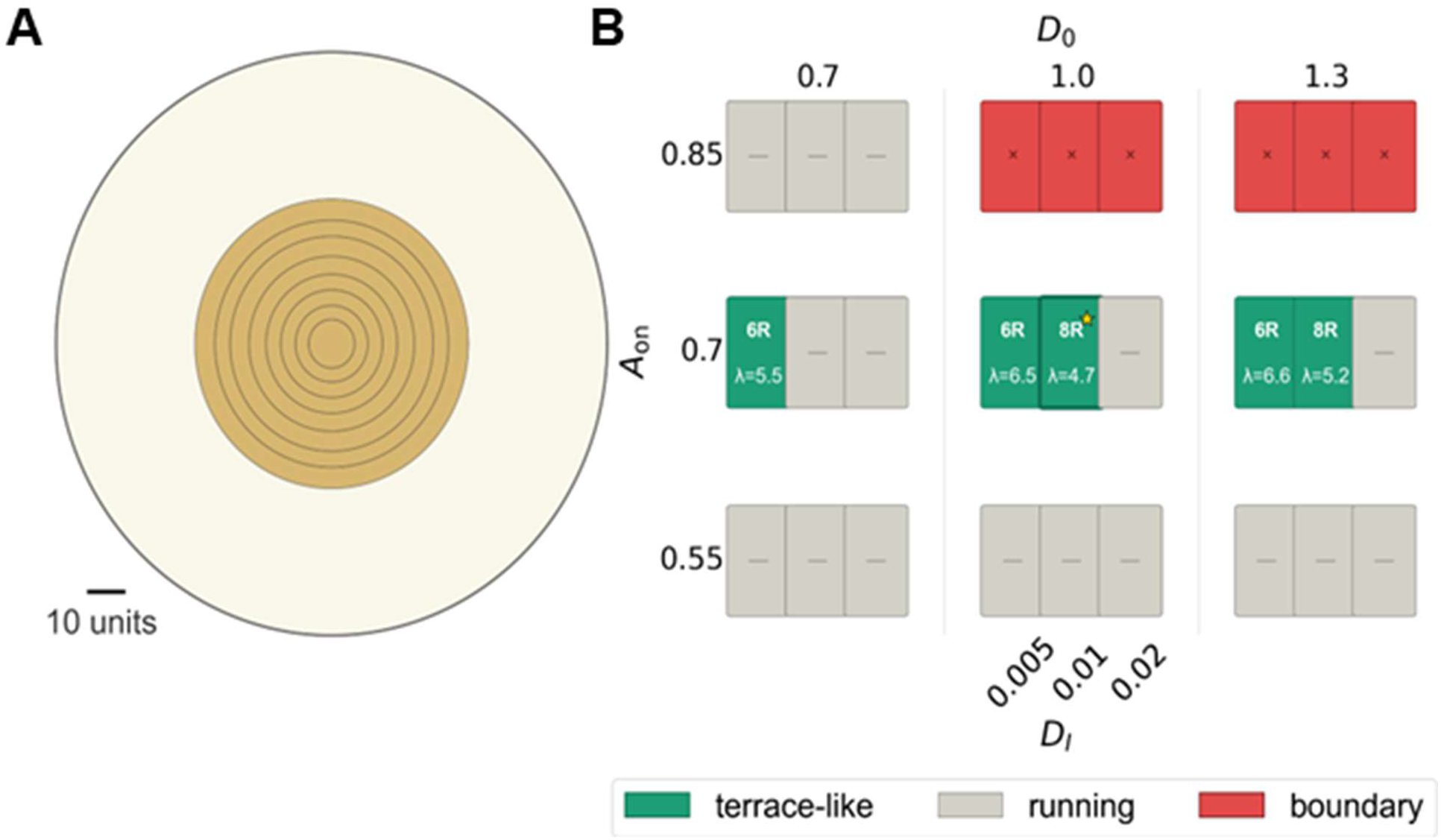
Concentric ring formation in the swarming–mineralization model. **(A)** Simulated ring pattern generated by rotating the one-dimensional radial solution *P*(*r*, *t*) around the inoculation centre under the assumption of perfect radial symmetry. (**B)** Parameter space survey over a 3×3×3 grid of motility coefficient *D_0_ ϵ* {0.70,1.00,1,30}, activation threshold *A_0n_ ϵ* {0.55,0.70,0.85} and inhibitor diffusivity *D_I_ ϵ* {0.005, 0.010, 0.020}. Green cells: ring formation regime; ring count (*nR*) and mean spacing (*λ̄*) are shown. The starred cell (*) indicates the representative case shown in Figure 4A.

To understand how rings form within this framework, we examined the space-time structure of the gate field *H*(*r*, *t*) and the temporal evolution of the front position *R*(*t*). The kymograph of *H*(*r*, *t*) reveals that each ring corresponds to a brief, spatially localized activation event near the advancing front (**Figure 5A**). Following each event, the site enters the refractory state and the region behind the front remains permanently inactive, confirming that each spatial location traverses the sequence READY → ACTIVE → REFRACTORY exactly once with no re-firing. Consistently, the front position R(t) follows a staircase trajectory in which each plateau corresponds to one arrest event and one deposited ring (**Figure 5B**).

**Figure 5.**
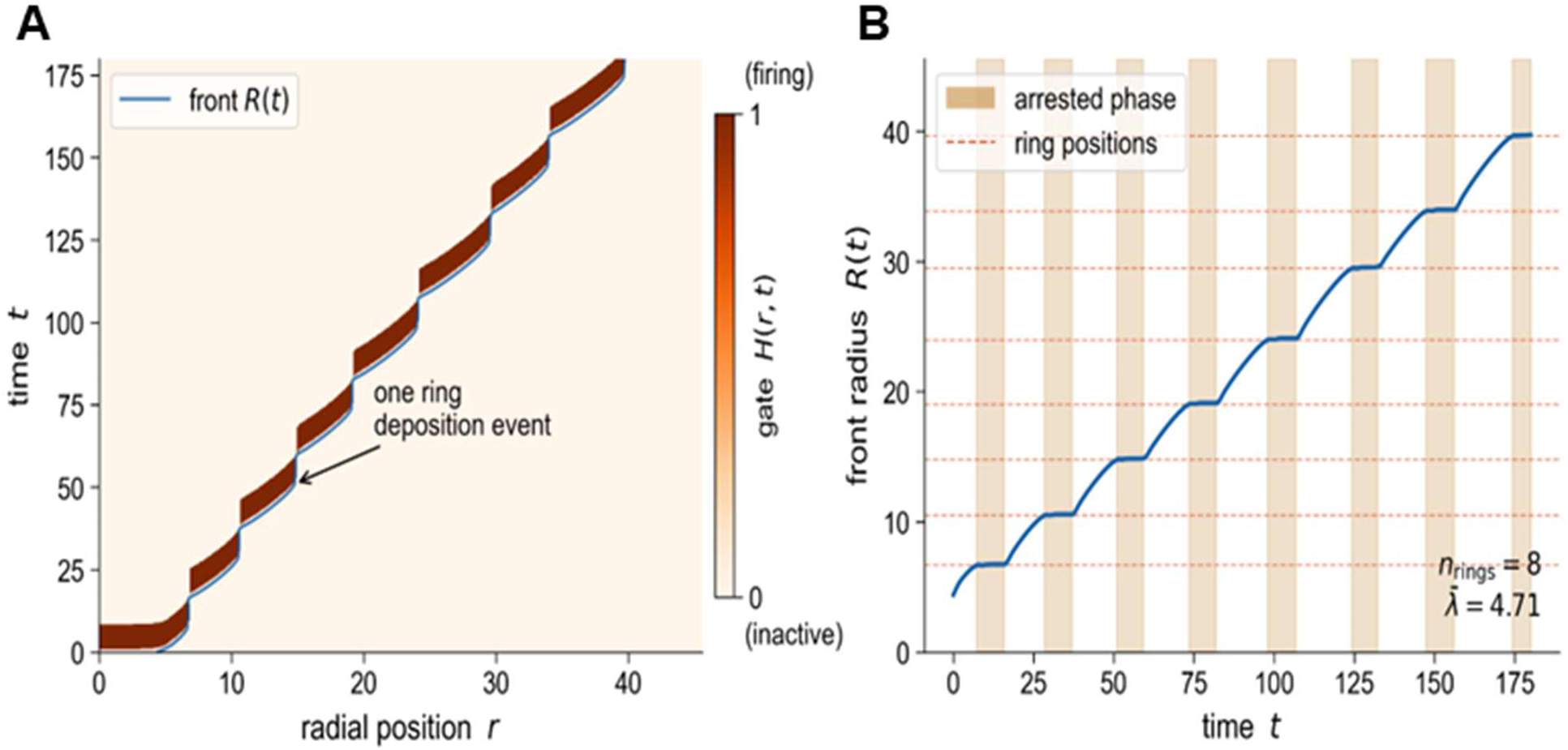
Space–time structure of ring formation. **(A)** Kymograph of the precipitation gate *H*(*r*, *t*). Eight narrow stripes localized near the advancing front (red curve) correspond to eight successive ring deposition events. **(B)** Front radius *R*(*t*) as a function of time. Dashed horizontal lines mark the eight ring positions inferred from front-velocity plateaus. The parameters are taken from **Table S1**.

The arrest-restart mechanism is dissected in **Figures 6A**–**6C**. Arrest is initiated when the inhibitor field *I* at the front rises above the *I_gate_* = 0.10 threshold (**Figure 6A**). This causes *D_eff_* to collapse by three to four orders of magnitude (**Figure 6B**), bringing the front velocity to zero (**Figure 6C**). Motion resumes only after the inhibitor decays below *I_gate_* and *D_eff_* recovers. The repeated cycle of inhibitor buildup, motility collapse, and inhibitor decay drives the periodic deposition of mineral rings. The bistable character of the local swarming dynamics that underpins this cycle is illustrated in **Figure S14**. Together, these results establish that concentric ring formation requires three cooperating elements: the bistable swarming-density field, the activation-inhibition cycle governed by local supersaturation, and irreversible refractory memory from mineral deposition.

**Figure 6.**
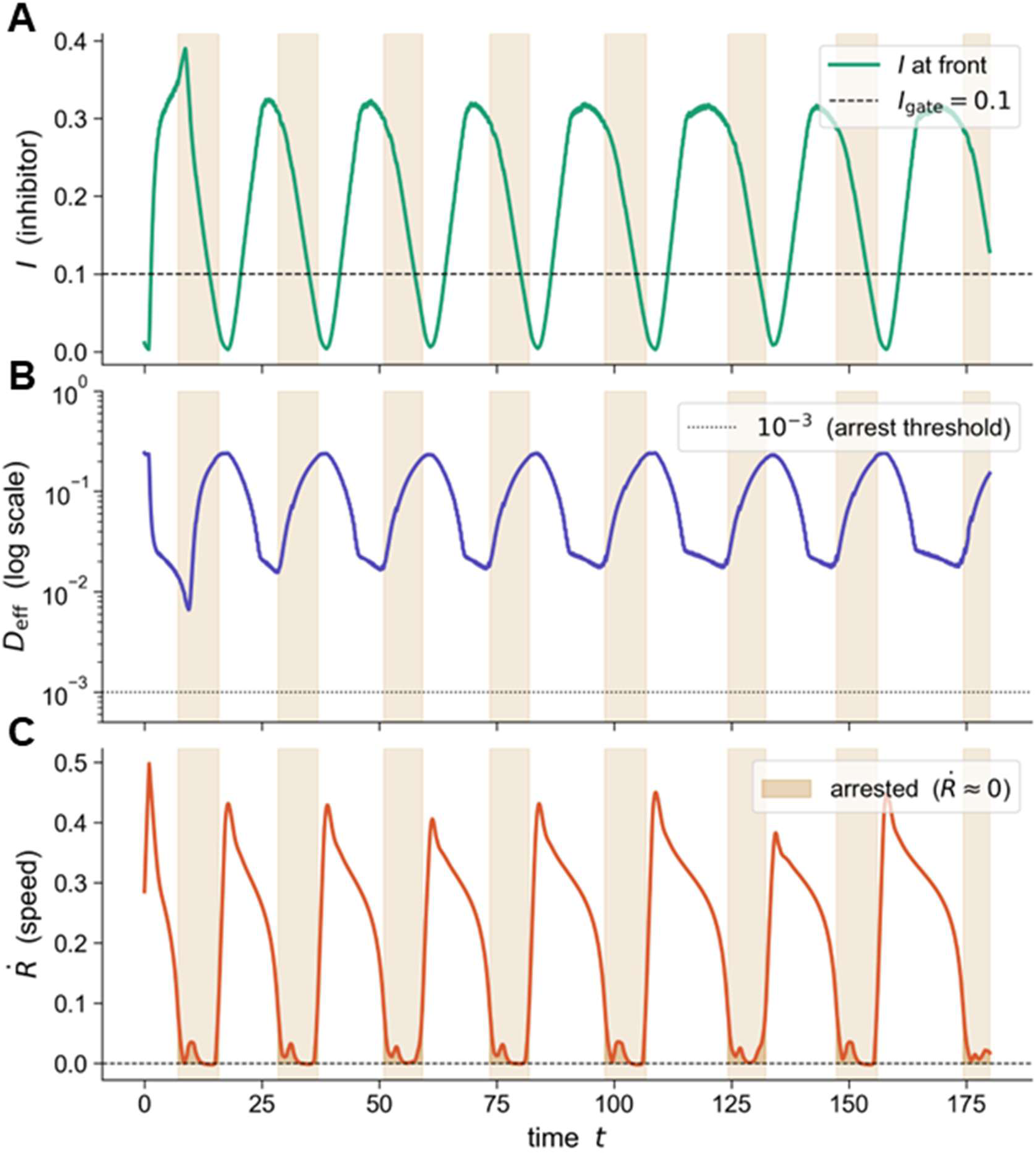
Mechanism of front arrest and restart. Orange bands indicate arrested phases as in Figure 5B. (A) Front-local inhibitor *I*(*r_front_*, *t*). **(B)** Front-local effective diffusivity *D_eff_*(*r_front_*, *t*) on a logarithmic scale. (C) Front speed *Ṙ*(*t*).

Having established that the model reproduces concentric ring formation mechanistically, we next asked how well its quantitative predictions align with experimental observations. The experimental ring positions *R*_1_ − *R*_6_were measured across three independent replicates (**Figure 7A**). The central inoculum circle was excluded as it reflects initial colony size rather than the ring location. The average experimental ring spacing was *λ̄_exp_* = 0.332 ± 0.024 mm. Because the model is dimensionless, we use normalized ring *λ_k_*/*λ̄* spacing to compare with experimental ring spacing (**Figure 7B**). Both experiment and simulation show a positive widening trend (experiment: +0.020 *λ̄*/ring, simulation: +0.088 *λ̄*/ring). Normalized spacings for ring pairs *R*_2_ → *R*_3_, *R*_3_ → *R*_4_, and *R*_4_ → *R_S_* agree within one standard deviation. We posit that the increase in ring space is due to the fact that, as the colony grows outward, the front biomass must cover a larger area. The bacteria have to spread over a larger area, which slows the leading edge and gives more time for arrest events to occur. Since ring spacing depends on both front speed and local generation time, slower front progression at larger radii naturally produces wider inter-ring distances. The quantitative discrepancy in the widening rate between simulation and experiment, most apparent at *R*_1_ − *R*_2_and *R_S_* − *R*_6_, likely reflects the model’s assumption of perfect radial symmetry and the omission of physical effects such as agar surface heterogeneity, nutrient gradients, and evaporation, which will be important to incorporate in future refinements of the model.

**Figure 7.**
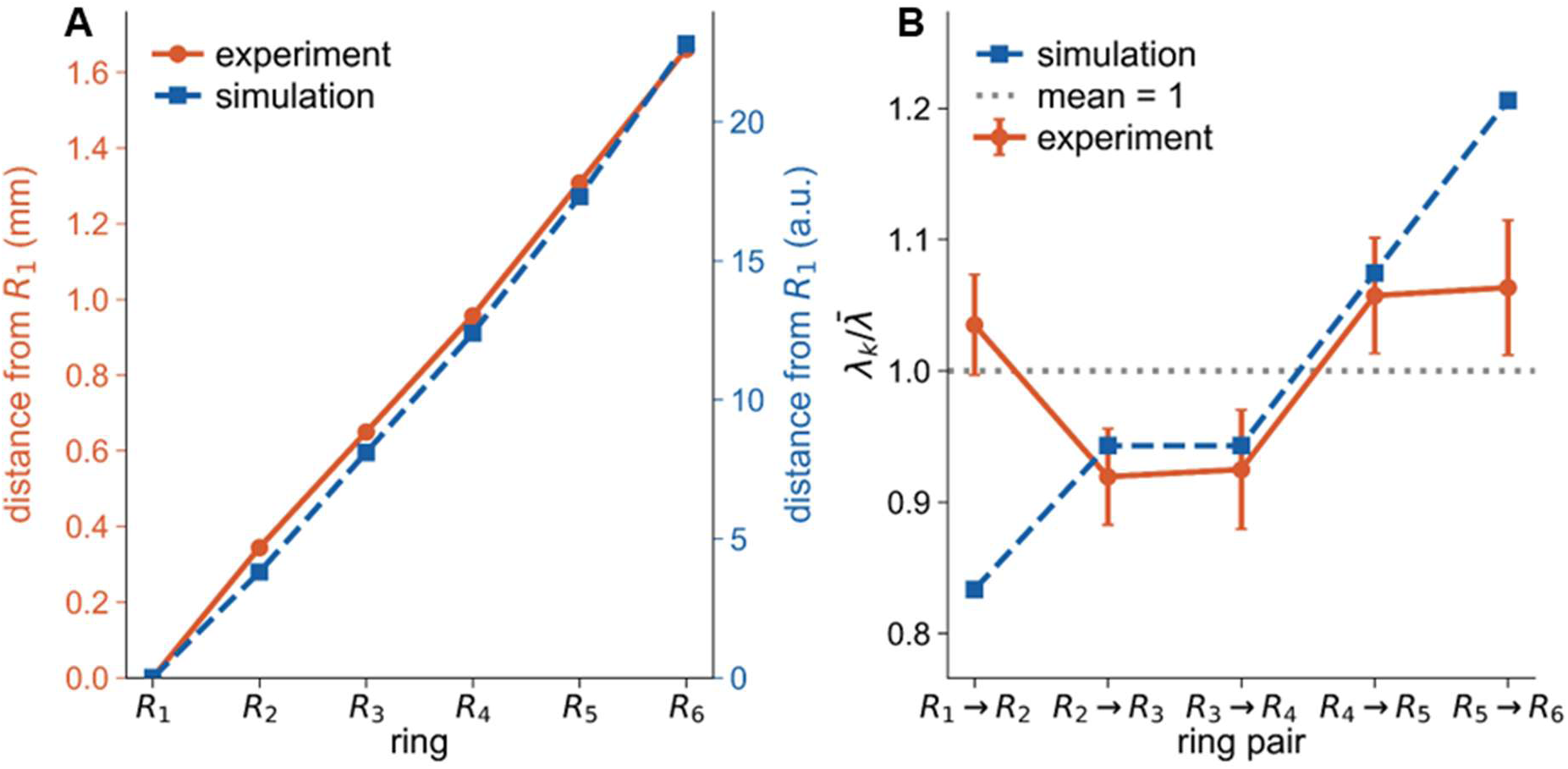
Ring spacing analysis comparing experimental measurements. Central inoculum circle excluded from analysis. **(A)** Cumulative ring positions measured from *R*1 for experiment (orange, mean ± SD, *n* = 3 replicates) and simulation (blue, right axis). **(B)** The ring spacing *λ_k_* is normalized by the mean spacing *λ̄* for each dataset independently. The reference line at *λ_k_*/*λ̄* = 1 corresponds to perfectly uniform spacing. Both experiment and simulation show a positive widening trend (experiment: +0.020 *λ̄*/ring, simulation: +0.088 *λ̄*/ring). Normalized spacings for ring pairs *R*_2_ → *R*_3_, *R*_3_ → *R*_4_, and *R*_4_ → *R_S_* agree within one standard deviation; deviations at *R*_1_ → *R*_2_ and *R_S_* → *R*_6_ reflect the inoculum transient and the growing radial, respectively.

## CONCLUSION

This study establishes bacterial swarming as a programmable platform for spatially organized biomineralization. Using E. coli MG1655, we show that phosphate sources alone exert negligible influence on motility, whereas calcium ions profoundly restrict swarming while simultaneously driving mineral deposition. Exploiting this interplay, we generated concentric mineralized rings that form and expand radially with the swarm front, a phenomenon not previously documented. SEM and EDS analyses confirm the deposits as calcium hydroxyapatite, with stoichiometry and microarchitecture sensitive to both calcium concentration and phosphate source identity. Environmental pH, modulated through glucose metabolism, further regulated the extent of both swarming and mineralization. A continuum model coupling density-dependent swarming with a mineralization-driven activation-inhibition cycle reproduced the observed ring patterns with minimal physical assumptions, identifying local supersaturation, irreversible mineral deposition, and motility suppression as the essential ingredients for periodic ring formation. Together, this work reframes bacterial swarming as a tunable materials synthesis process in which collective microbial behavior encodes spatial information directly into mineral structure, opening avenues for engineering patterned biomineralization without scaffolds or external templating, with prospective applications in biomedicine, materials science, and bio-inspired surface design.

## AUTHOR CONTRIBUTIONS

K.P. conceived the original idea and planned the experiments. K.V.C. planned and carried out experiments and data analysis. U.K. carried out the modelling and relevant computational work.

K.P. supervised the research along with providing feedback during the writing of the manuscript.

All authors provided critical feedback and helped shape the research, analysis, and manuscript.

## ACKNOWLEDGEMENTS

We thank Prof. Chandan Mishra for reading and providing critical feedback on this work. The authors thank CRTDH (Common Resource and Technology Development Hub) and CIF (Central Instrumentation Facility) for providing various instrumentation facilities at IIT Gandhinagar.

