## Supplementary figures and Table for "Bacterial Swarming-Guided Biomineralization Enables Pattern Formation in Engineered Living Materials"

*To whom all correspondence must be addressed

Karthik Pushpavanam, Ph.D.

Department of Chemical Engineering

Indian Institute of Technology Gandhinagar

Gujarat, India, 382055, India

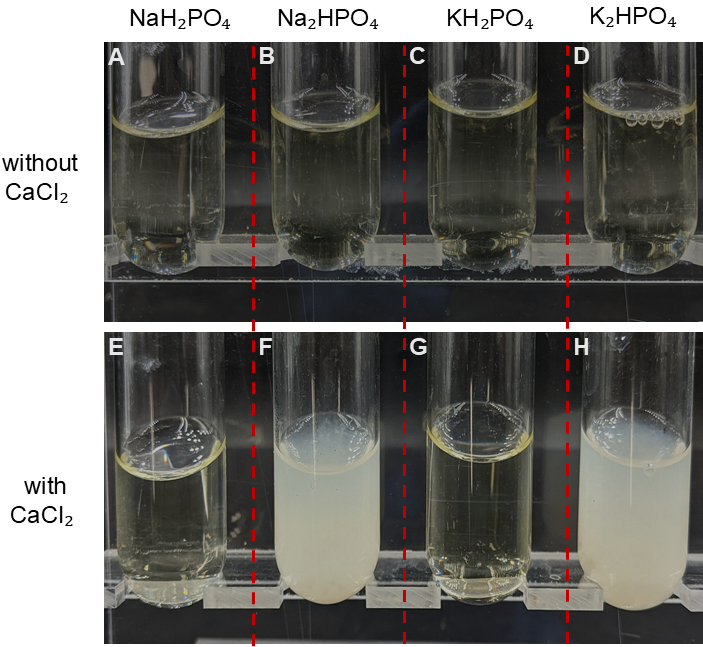

**Figure S1. Differential Calcium precipitation in the presence of monobasic- and dibasic phosphates.** The precipitation of calcium as cloudy precipitates in Nutrient broth (**A-D**) without 10mM calcium chloride and (**E-H**) with 10mM calcium chloride in the presence of NaH₂PO₄, Na₂HPO₄, KH₂PO₄, and K₂HPO₄ at 10mM concentration.

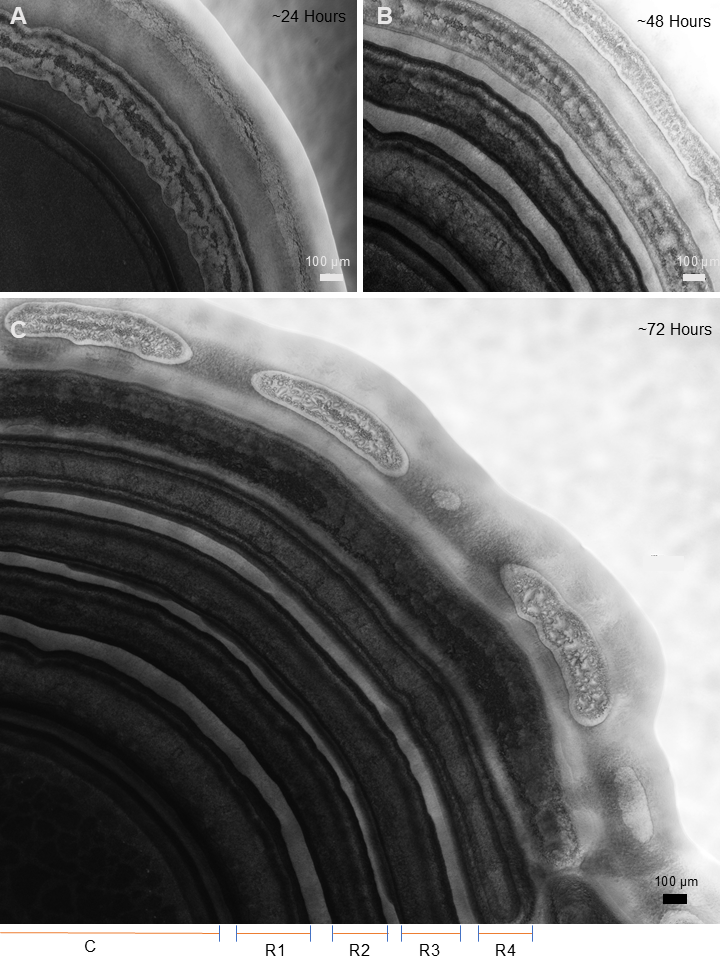

**Figure S2. Variation of the distance between two consecutive concentric rings within a swarm.** Phase-contrast image of discontinuous calcium mineralization of swarming bacteria observed after (**A)** 24, **(B)** 48, and **(C)** 72 hours of incubation at 37℃ on nutrient agar plate supplemented with 10 mM NaH₂PO₄ and 10 mM CaCl₂, with 5 μl of volume spotted. The different zones of the mineralization pattern, namely the central region (C), Ring-1 (R1), Ring-2 (R2), Ring-3 (R3), and Ring-4 (R4), are shown with their approximate boundaries.

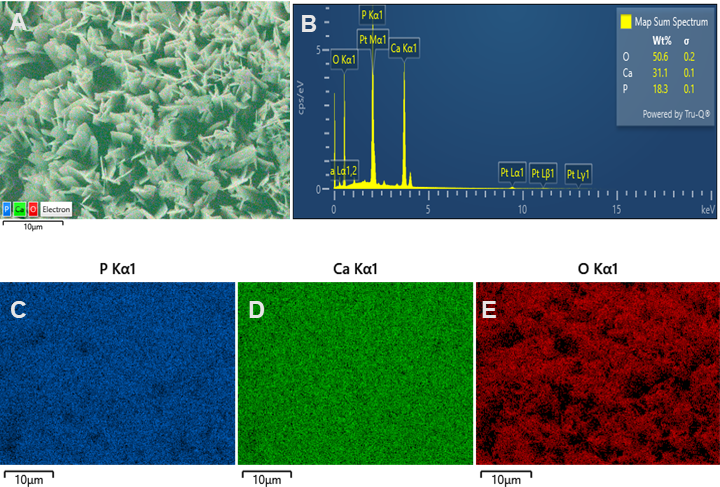

**Figure S3. EDS elemental composition of swarm grown in 10 mM NaH₂PO₄ and 10mM CaCl₂.** The elemental composition of a region on the dried swarm surface with **(A)** EDS layered image, **(B)** corresponding weight percentage map of elements, and elemental density maps representing (**C)** phosphorus, **(D)** calcium, and **(E)** oxygen.

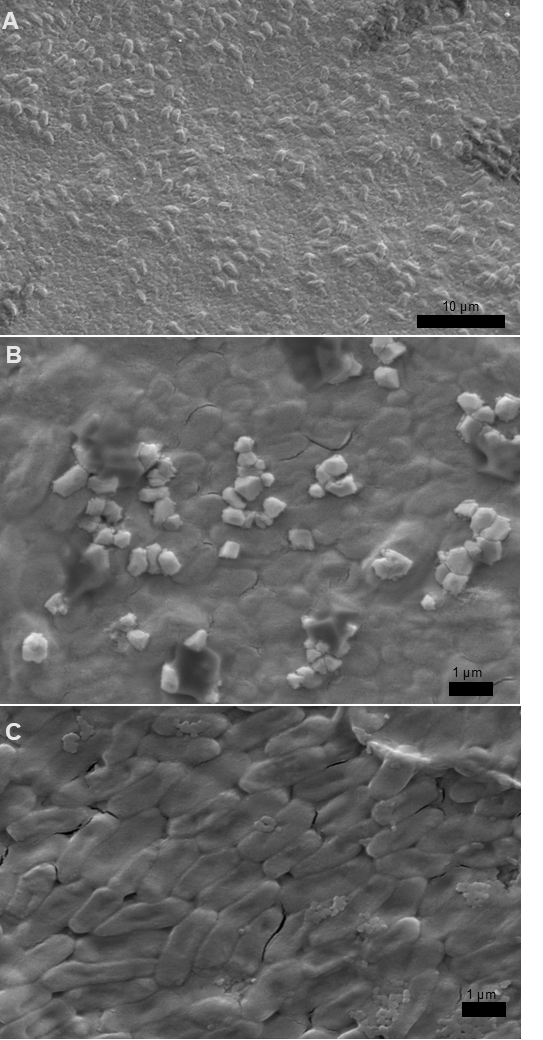

**Figure S4. The difference between bacterial swarms grown in different conditions.** SEM images of bacteria swarm grown in (**A**) Nutrient Broth 03 (control), (**B**) 10mM NaH₂PO₄, and (**C**) 5mM calcium chloride.

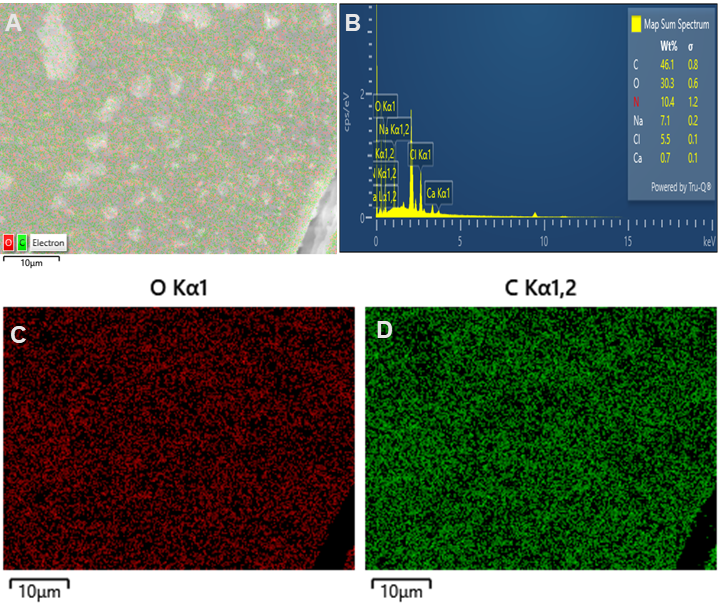

**Figure S5. EDS elemental composition of the swarm grown in the nutrient broth 03 (control).** The elemental composition of a selected area on the dried swarm surface with **(A)** EDS layered image, **(B)** corresponding weight percentage map of elements, and elemental density maps representing (**C)** oxygen, and **(D)** carbon.

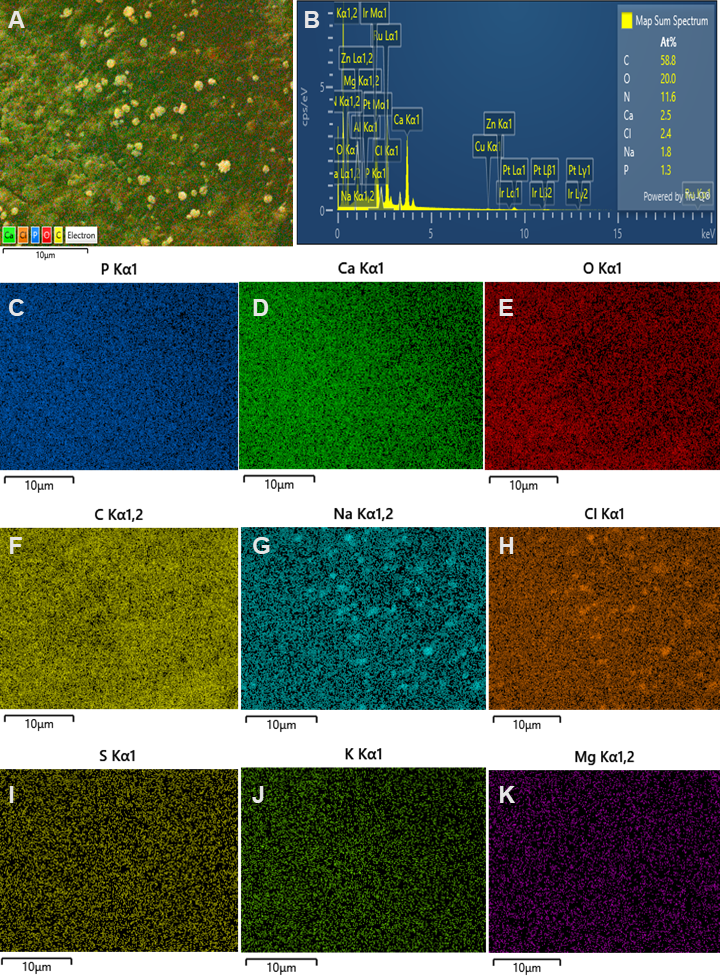

**Figure S6. EDS elemental composition of swarm grown in 5mM CaCl₂.** The elemental composition of the selected region on dried swarm surface with **(A)** EDS layered image **(B)** corresponding weight percentage map of elements, and elemental density maps representing **(C)** phosphorus, **(D)** calcium, **(E)** oxygen, **(F)** carbon, **(G)** sodium, **(H)** chlorine, **(I)** sulphur, **(J)** potassium, and **(K)** magnesium

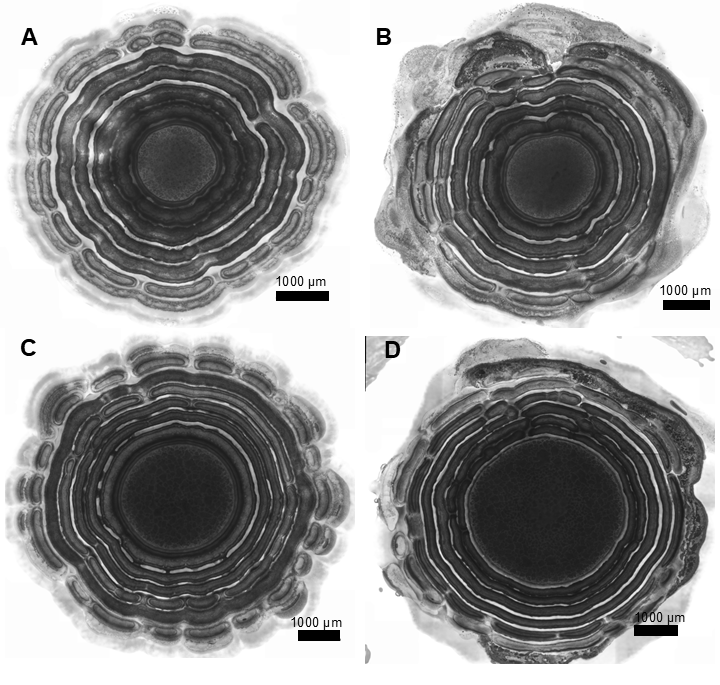

**Figure S7. Effect of inoculum volume on swarming patterns.** Phage-contrast images of calcium mineralization of swarming bacteria observed after incubation for 4 days at 37℃ on nutrient agar plate supplemented with 10 mM NaH₂PO and 5mM CaCl₂, spotted using **(A)** 1.25 μl, **(B)** 2.5 μl**, (C)** 5 μl and **(D)** 10 μl of the bacterial culture.

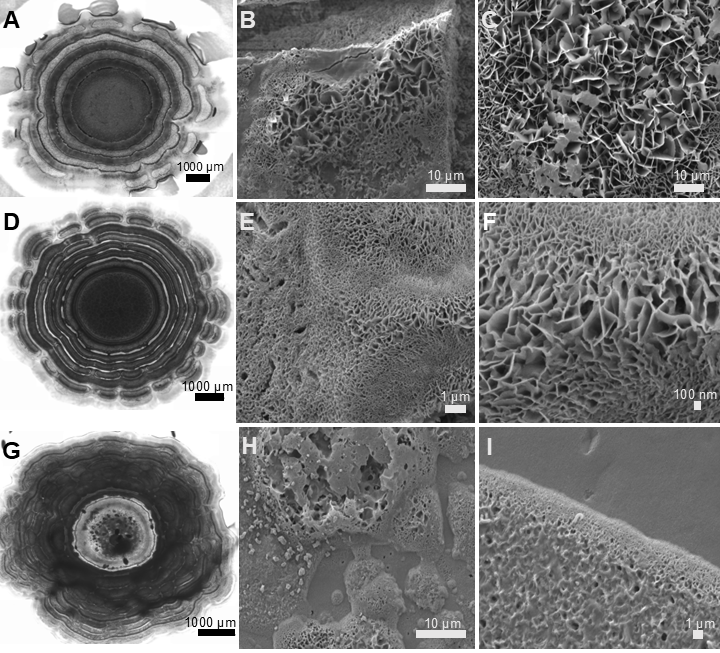

**Figure S8. Effect of calcium concentration on mineralizing bacterial swarm.** Phase-contrast and corresponding SEM images of calcium mineralization of swarming bacteria observed after incubation for 4 days at 37℃ on nutrient agar plate supplemented with 10 mM NaH₂PO₄, along with **(A, B, C)** 5mM CaCl₂, **(D, E, F)** 10 mM CaCl₂, and **(G, H, I)** 20 mM CaCl₂, with 5 μl of volume spotted.

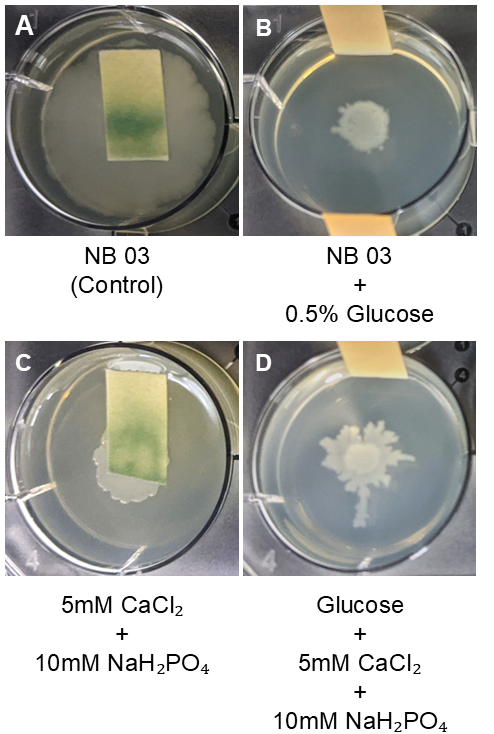

**Figure S9. Acidification of media via glucose metabolism.** The pH indicator strips showing basic (green) and acidic (orange) in the swarming media after incubation for 4 days at 37℃ grown in **(A)** Nutrient broth 03 (control), along with **(B)** with 10mM CaCl₂ and 10 mM NaH₂PO₄, **(C)** 0.5% glucose, and **(D)** 0.5% glucose added with 5mM CaCl₂ and 10 mM NaH₂PO₄.

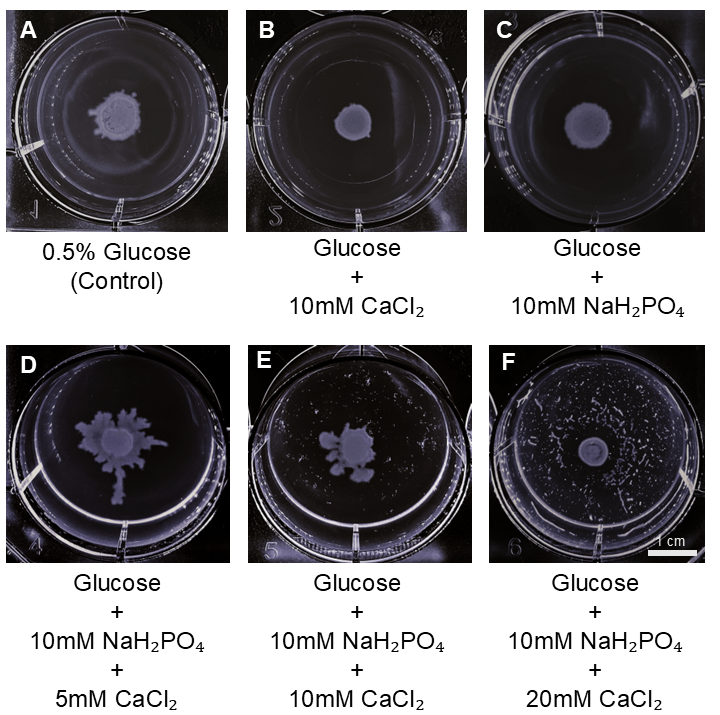

**Figure S10. Acidic pH negatively affects mineralization during bacterial swarming.** The differences in the swarming pattern of bacteria observed after incubation for 4 days at 37℃ grown in glucose supplemented Nutrient broth with **(A)** 0.5% glucose (control), **(B)** 10mM CaCl₂, **(C)** 10 mM NaH₂PO₄, **(D)** 10 mM NaH₂PO₄ and 5mM CaCl₂, **(E)** 10 mM NaH₂PO₄ and 10mM CaCl₂, and, **(F)** 10 mM NaH₂PO₄ and 20mM CaCl₂.

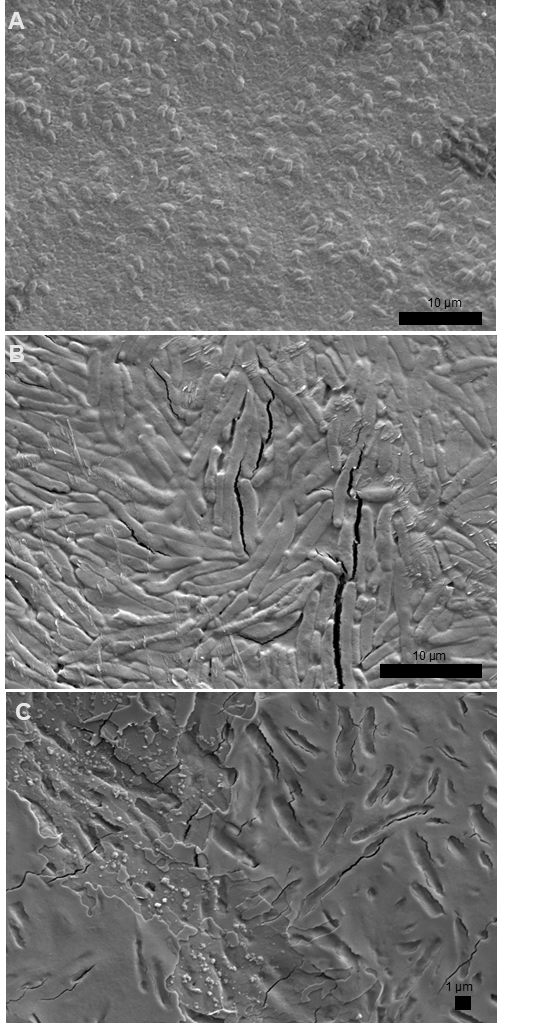

**Figure S11. The difference between bacterial swarms grown in different conditions.** SEM images of bacteria swarm grown in (**A**) Nutrient Broth 03 (control), (**B**) 0.5% Glucose, and (C) 0.5% Glucose with 10mM calcium chloride and 10mM NaH₂PO₄.

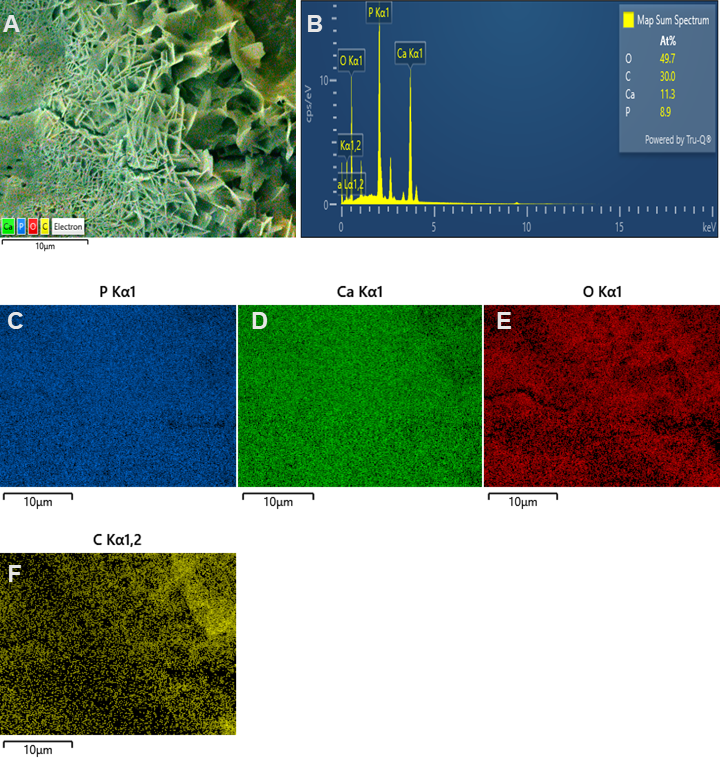

**Figure S12. EDS elemental composition of swarm grown in 10mM CaCl₂ and 10mM KH₂PO₄.** The elemental composition of the selected region on the dried swarm surface with (**A)** EDS layered image **(B)** corresponding weight percentage map of elements, and elemental density maps representing (**C)** phosphorus, **(D)** calcium, (**E)** oxygen, and **(F)** carbon.

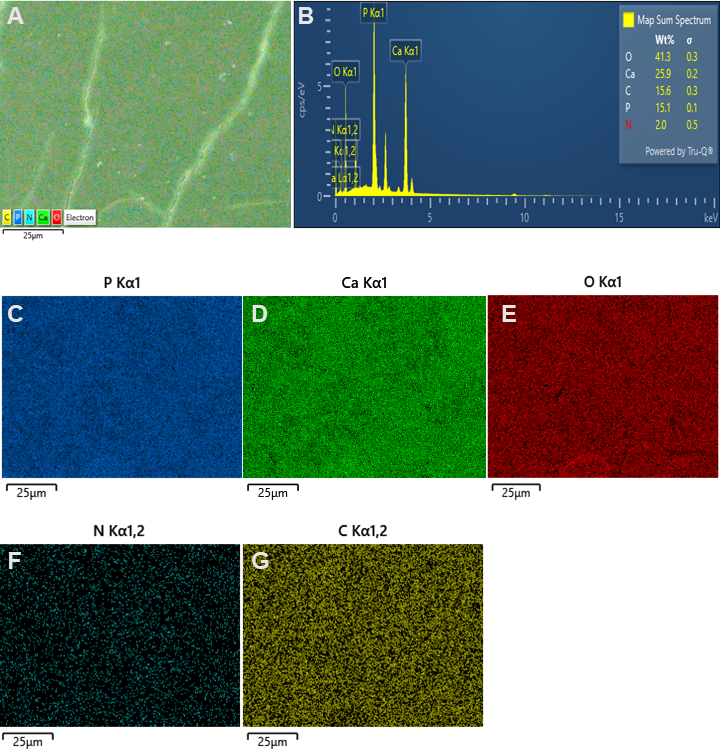

**Figure S13. EDS elemental composition of swarm grown in 10mM CaCl₂ and 10mM β-Glycerophosphate.** The elemental composition of the selected region on the dried swarm surface with (**A)** EDS layered image, **(B)** corresponding weight percentage map of elements, and elemental density maps representing (**C)** phosphorus, **(D)** calcium, **(E)** nitrogen, and **(F)** carbon.

**BISTABLE SWARMING KINETICS**

To analyze the bistable nature of the swarming density field, we consider a well-mixed location with no spatial variation and inhibitor coupling. We reduce Eq. (1) by setting $D_{eff}=0, s0=0, I=0$.

The resulting ordinary differential equation

$$\frac{dP}{dt}=f(P)-\mu P,$$

(S1)

where,

$$f\left( p \right)=\frac{\alpha P^{n}}{K_{p}^{n}+P^{n}} \left( 1-\frac{P}{P_{max}} \right).$$

(S2)

The system has three fixed points: the trivial state $P=0$ , an unstable state at $P_{unstable}^{*}\approx0.170$, and a stable state at $P_{wake}^{*}=0.631$.

The unstable fixed-point acts as a threshold. Below this, local swarming activity decays, and above this, it grows toward a stable high state. This stable high state represents the colonized region behind the moving front. Because this value remains far above the motility threshold $P_{min} = 0.05$, the colony interior remains active even during arrest phases. This allows the front to restart once inhibition weakens. The front-detection threshold $(P_{thresh}= 0.22)$ is chosen within this growing regime to track the front consistently. Parameter values are given in **Table S1**.

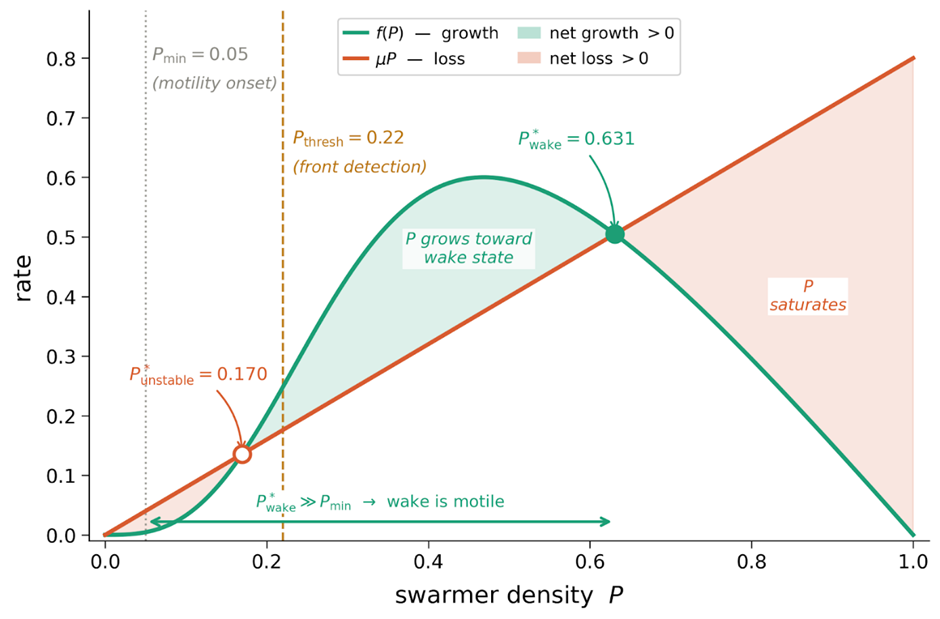

**Figure S14**. **Bistable kinetics of the swarming density field**. Bistable kinetics of the swarmer density field $P$, obtained from the spatially homogeneous limit of Eq. (1) with diffusion, inhibitor, and source terms set to zero. The nonlinear growth rate $f(P)$ and linear loss rate $\mu P$ intersect at an unstable fixed point $(P_{unstable}^{*}=0.170)$ and a stable wake state $(P_{wake}^{*}=0.631)$. Since $P_{wake}^{*}\gg P_{min}=0.05$, the colony interior remains motile after each arrest event. The front detection threshold $P_{thresh}=0.22$ lies within the net-growth region, between the two fixed points.

**THRESHOLD PARAMETER CALCULATION**

The threshold parameters $A_{on}$, $A_{off}$, $P_{gate}$, and $I_{gate}$ control the state transitions and are chosen from the consistency conditions set by the model’s steady states. The front-activity threshold $P_{gate}$ is placed between the motility threshold $P_{min}$ and the unstable fixed point $P_{unstable}^{*}$ of the bistable *P* dynamics (**Figure S14**), so that a deposition event begins only when the swarming front is genuinely present.

During the running phase ($H=0$), setting $\partial_{t}A=0$ in Eq. (2) gives the resting steady state

$$A_{rest} =\frac{k_{form}}{k_{loss}} \Phi(P).$$

(S3)

The activation threshold $A_{on}$ is chosen below $A_{rest}$, which allows the activation field to build up during the active phase.

During active deposition ($H=1$), setting $\partial_{t}A=0$ gives the active steady state

$$A_{active}=\left[ \frac{k_{form}-k_{cons}}{k_{loss}} \right]\Phi(P).$$

(S4)

The threshold $A_{off}$ is set above $A_{active}^{*}$ , so that each firing event ends when $A$ drops below $A_{off}$. Therefore, the thresholds must satisfy

$$A_{active}< A_{off}<A_{on}<A_{\mathrm{rest}}.$$

(S5)

During active deposition ($H=1$), setting $\partial_{t}I=0$ in Eq. (3) and ignoring diffusion gives the maximum inhibitor level

$$I_{peak}=\alpha_{I}\tau_{I}\Phi\left( P^{*} \right).$$

(S6)

The inhibitor gate $I_{gate}$ is chosen below $I_{peak}$, so that a new deposition event cannot begin until the inhibitor has decayed sufficiently.

After the deposition event ends ($H=0$), $I$ decays as

$$I(t) = I_{peak}exp(-t / \tau_{I})) .$$

(S7)

This sets $I\left( T_{off} \right)=I_{gate}$ and gives the recovery duration

$$T_{off}= \tau_{I} ln(I_{peak}/I_{gate}),$$

(S8)

which connects the gate value directly to the observable inter-event recovery time.

**Table S1: Model parameters used in the simulations**

| **Symbol** | **Description** | **Value** |
| --- | --- | --- |
| $D_{0}$ | Base motility scale for effective bacterial swarming | $1$ |
| $P_{min}$ | Motility onset threshold for coordinated swarming | $0.05$ |
| $\beta_{I}$ | Strength of motility suppression by mineralization inhibition | $12$ |
| $s0$ | Basal source term for swarming | $4\times{10}^{-3}$ |
| $\alpha$ | Nonlinear activation strength of swarming | $1.6$ |
| $K_{p}$ | Half-saturation constant for swarmer activation | $0.35$ |
| $n$ | Hill exponent | $3$ |
| $P_{max}$ | Saturation level of swarmer activity $P$ | $1$ |
| $\mu$ | 𝐷ecay rate of swarmer activity $P$ | $0.8$ |
| $P_{sat}$ | Saturation scale in $\Phi\left( P \right)$ | $0.45$ |
| $D_{A}$ | Diffusivity of the local activation state $A$ | $0$ |
| $k_{form}$ | Buildup rate of the local activation state $A$ | $3.0$ |
| $k_{loss}$ | Relaxation rate of the local activation state $A$ | $0.45$ |
| $k_{cons}$ | Consumption rate of A during an activation even | $2.9$ |
| $A_{on}$ | Threshold for transition from READY to ACTIVE | $0.70$ |
| $A_{off}$ | Threshold for transition from ACTIVE to REFRACTORY | $0.08$ |
| $D_{I}$ | Diffusivity of mineralization-related inhibition $I$ | $0.010$ |
| $\alpha_{I}$ | Production strength of mineralization-related inhibition | $4$ |
| $\tau_{I}$ | Relaxation time of mineralization-related inhibition | $0.8$ |
| $P_{gate}$ | Front swarming threshold required for site activation | $0.08$ |
| $I_{gate}$ | Maximum inhibition level allowing activation | $0.10$ |
| $L$ | Domain radius | $80$ |
| $N_{r}$ | Spatial grid points | $900$ |
| $dt$ | Time step used in RK4 integration | $5\times{10}^{-4}$ |
| $T$ | Total simulation time | $180$ |
